# The cornerstone of early diagnosis and immunotherapy of prostate cancer:screening characteristic genes

**DOI:** 10.1101/2024.05.15.594356

**Authors:** Bo Shao, Kaixiu Wu, Shui Wan, Pingping Sun, Yanggen Zuo, Li Xiao, Jinbo Pi, Zhengkai Fan, Zhongxiong Han

## Abstract

**Background:** Prostate cancer (PCA) has become a common malignant tumor globally, posing a substantial risk to the health of middle-aged and elderly men. However, there is still a lack of effective strategies for early detection and treatment of prostate cancer. The introduction of gene therapy in recent years has shown promise as a potential approach for cancer diagnosis and treatment.

**Methodology & Theoretical Orientation:** The training set data GSE45016, GSE46602, and GSE69223 from the Gene Expression Omnibus (GEO) dataset, along with validation training set data GSE17951, were utilized. Differentially expressed genes (DEGs) between normal individuals and tumor patients were identified by combining the training set data. Subsequent analyses including Gene Ontology (GO), Kyoto Encyclopedia of Genes and Genomes (KEGG), and gene set enrichment analysis (GSEA) were conducted on the DEGs. WGCNA analysis was then performed on the gene expression matrix to identify module genes highly correlated with PCA, followed by the application of the LASSO algorithm to obtain PCA candidate genes. The candidate genes were validated using the area under the receiver operating characteristic (ROC) curve (AUC) to determine key feature genes. Finally, the relationship between key characteristic genes and immune cells was explored.

**Findings:** A total of 54 DEGs were identified, with 26 down-regulated genes and 28 up-regulated genes. The GO function analysis revealed enrichment in processes such as ‘establishment of protein localization to membrane’ and ‘protein targeting to membrane’. KEGG analysis showed enrichment in pathways like ‘eutrophil degranulation’, ‘neutrophil activation involved in immune response’, and ‘regulation of cell morphogenesis’. GSEA analysis highlighted enrichment in pathways like ‘CTRL_VS_ACT_IL4 AND ANTI_IL12_12H_CD4_TCELL_DN’. Through WGCNA and LASSO regression analysis, key characteristic genes MARCKSL1, TMTC4, and TTLL12 were identified, with AUC values greater than 0.8 in both the training and validation sets, and were found to be closely associated with immune cell infiltration.

**Conclusion & Significance:** MARCKSL1, TMTC4, and TTLL12 emerge as crucial genes in the process of PCA, showing significant relevance to immune cell infiltration.this study offers valuable clinical insights into the diagnosis and treatment of prostate cancer through the identification of specific genes associated with the disease.

## 1 INTRODUCTION

Prostate cancer (PCA) is a prevalent malignant tumor of the urinary system that poses a significant threat to the health and lives of middle-aged and elderly men globally. In 2020, global cancer statistics revealed that PCA ranked second in male morbidity and sixth in mortality worldwide, with a morbidity rate of 14.1% and a mortality rate of 6.8%^[1]^. According to the American Cancer Society’s statistical analysis for 2023, PCA was the most common male tumor in the United States, with a morbidity rate of 29% and a mortality rate of 11%^[2]^. The prognosis of PCA is heavily influenced by the tumor grade and stage at the time of diagnosis: patients with early or localized PCA have a 10-year survival rate exceeding 99%, whereas those with advanced or metastatic disease at diagnosis have a 5-year survival rate of only 30%^[3]^. Early detection of PCA is crucial for patient prognosis, and studying key characteristic genes involved in the development of PCA can aid in early diagnosis and treatment.

Currently, disease risk prediction through bioinformatics research has emerged as a prominent area of study^[4-5]^. The advancement of bioinformatics has facilitated the identification of disease-specific genes using public data, significantly enhancing the accuracy of disease risk prediction. Given the current absence of key PCA-specific genes for early clinical diagnosis, we utilized bioinformatics methods to identify crucial genes associated with PCA. This research aims to support early prevention, diagnosis, treatment, and ultimately, improve the survival rate of patients with PCA.

## 2 MATERIAL AND METHODS

### 2.1 Data source

Four PCA-related microarray datasets were obtained from the Gene Expression Omnibus (GEO) (https://www.ncbi.nlm.nih.gov/geo/). GSE45016, GSE46602, and GSE69223 were utilized as training sets, while GSE17951 was employed as the validation set.

### 2.2 Research methods

#### 2.2.1 Data merging

The training set data was merged using the R language with the ‘limma’ package and ‘sva’ package. Differential expression analysis was then conducted on the merged data (|logFC| > 2 and P< 0.05), followed by the generation of heat maps and volcano plots.

#### 2.2.2 GO, KEGG and GSEA analysis

The differentially expressed genes (DEGs) were analyzed using the R programming language with packages such as limma, DOSE, clusterProfiler, and enrichplot. Gene Set Enrichment Analysis (GSEA) was conducted on the integrated data.

#### 2.2.3 Machine learning algorithm screening PCA candidate genes

Weighted gene co-expression network analysis (WGCNA) was conducted on the integrated gene expression matrix using the R software ‘WGCNA’ package. Initially, the weight parameter (power) value of the adjacency matrix was determined to construct the TOM matrix gene clustering tree based on this power value, establishing the weighted co-expression network model. The integrated gene expression matrix was then partitioned into related modules, and the module genes showing the highest correlation with PCA were selected for further analysis. The ‘veen’ package in R software was utilized to intersect the selected differential genes with the related genes identified through WGCNA analysis. The overlapping genes were subjected to LASSO analysis to identify candidate genes for PCA.

#### 2.2.4 Validation of key characteristic genes and their immune analysis

The R software package was utilized to analyze differential expression levels of candidate genes in normal and tumor tissues using the ‘ggpubr’ package in both the training and test sets. Receiver operating characteristic (ROC) curve analysis was conducted on both sets, identifying genes with area under curve (AUC) values exceeding 0.80 as key characteristic genes of PCA. Additionally, the correlation between these key genes and immune cells in PCA was examined using the ‘reshape2’, ‘ggpubr’, and ‘ggExtra’ packages.

#### 2.2.5 Statistical Methods

Data analysis and processing were carried out using R x64 version 4.0.2, with statistical significance set at P<0.05.

## 3 RESULTS

### 3.1 Screening and analysis of DEG

By combining the training set data with the differential analysis, DEG heat map (Figure 1A) and volcano map (Figure 1B) were generated, resulting in a total of 54 differential genes, with 26 down-regulated genes and 28 up-regulated genes identified. Enrichment analysis revealed that the biological process (BP) in the GO function of DEG was primarily associated with ‘establishment of protein localization to membrane’,The cellular component (CC) was predominantly found in ‘focal adhesion’ and ‘cell-substrate junction’,Molecular function (MF) was enriched in ‘actin binding’ (Fig. 1C).The KEGG analysis of DEG highlighted pathways such as ‘neutrophil degranulation’, ‘neutrophil activation involved in immune response’, and ‘neutrophil activation’(Fig. 1D).GSEA analysis highlighted enrichment in pathways like ‘CTRL_VS_ACT_IL4 AND ANTI_IL12_12H_CD4_TCELL_DN’(Fig. 1E,F).

**Figure 1:**
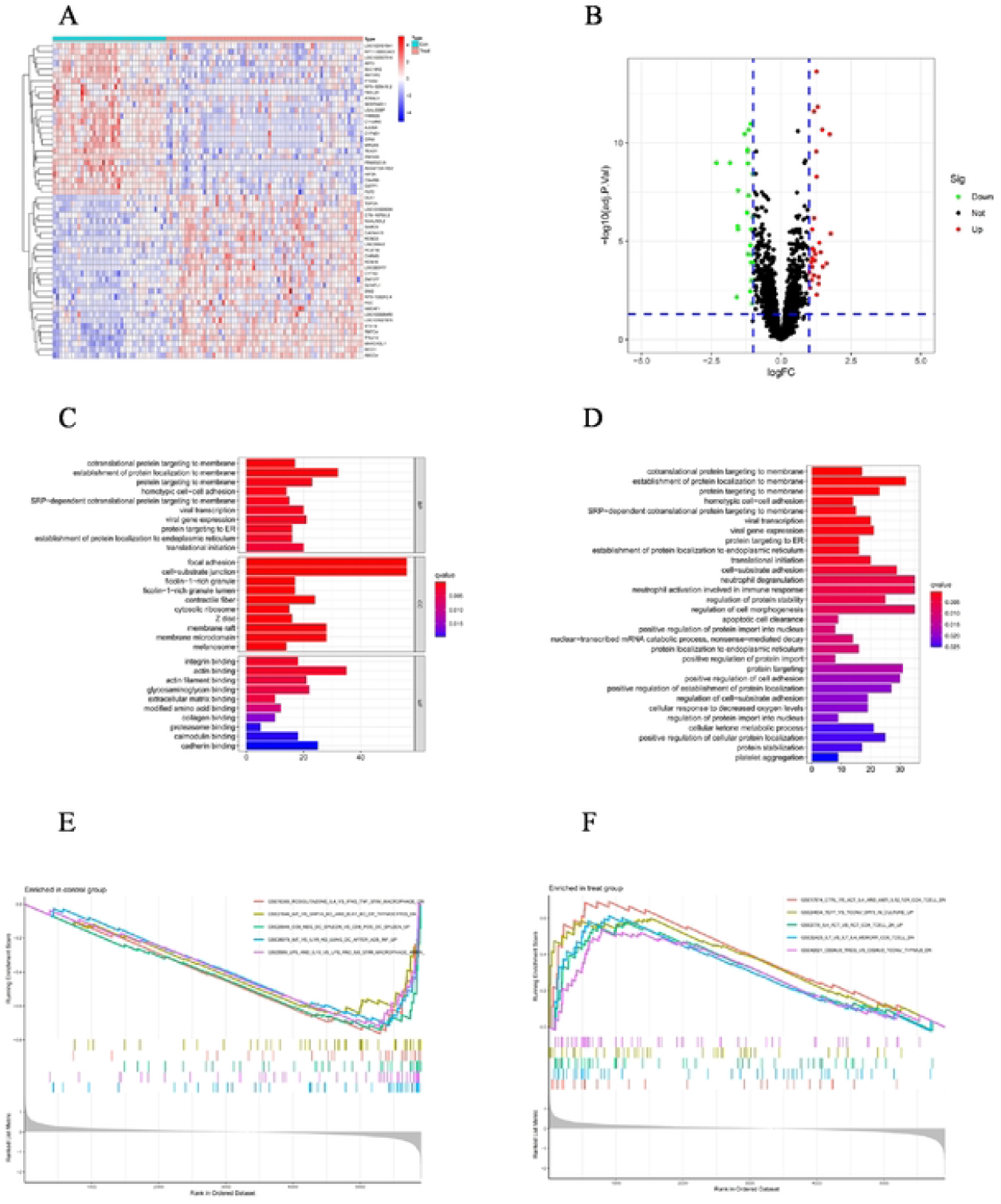
Figures A and B display genes with significant differences in expression between normal individuals and patients with PCA. In Figure A, ‘Con’ denotes normal individuals while ‘Treat’ represents PCA patients. Red signifies high expression levels and blue signifies low expression levels. In Figure B, red indicates up-regulated genes in tumor tissues, green represents down-regulated genes, and black denotes genes with no significant change in expression.Figure C shows the results of GO analysis, and Figure D shows the results of KEGG analysis. The redder the color, the more significant the pathway of differential gene enrichment.Figure E and Figure F represent the results of GSEA analysis of genes in normal people and PCA patients, respectively.

### 3.2 Machine learning algorithm screening PCA candidate genes

In this study, the weight parameter of the adjacency matrix was set to a power value of 10, resulting in a higher gene connection degree and a network structure closer to a scale-free network. Utilizing the weighted correlation coefficient of this power value, gene clustering in the TOM matrix was performed (see Figure 2A). Subsequently, a weighted co-expression network model was created to partition the gene expression matrix into four modules. The Pearson algorithm was then applied to calculate the correlation coefficient between the characteristic genes of each module and the traits. Notably, the turquoise color module exhibited a correlation coefficient of 0.76, P = 3e-20, with the highest correlation observed with PCA (see Fig. 2B). A total of 727 module genes were identified, and further filtering criteria were applied: geneSigFilter > 0.5 and moduleSigFilter > 0.75, resulting in the selection of 11 module genes(Figure 2C).

**Figure 2:**
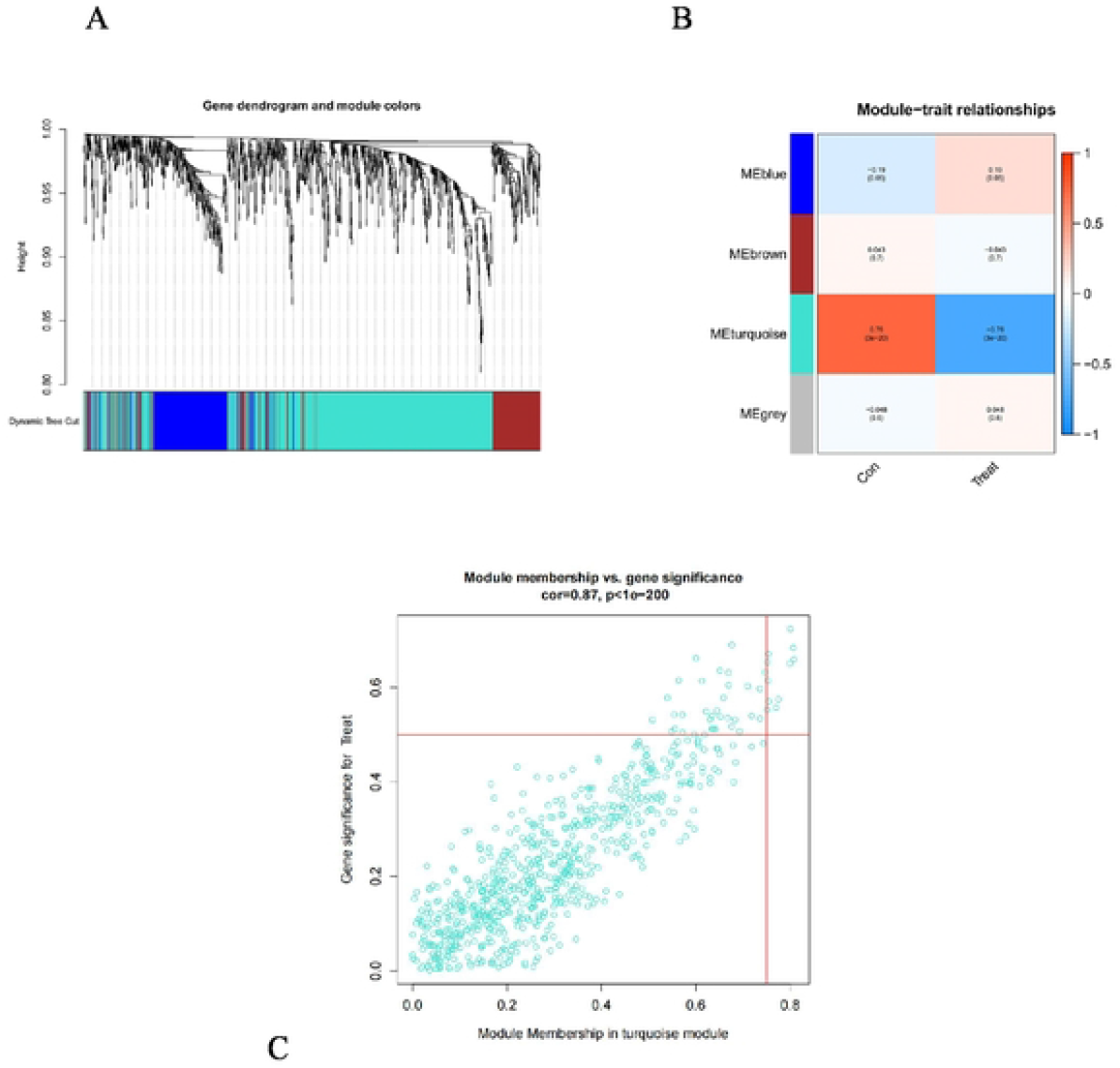
The results include TOM matrix gene clustering results (Fig. 2A), correlation coefficients between characteristic genes and traits (Fig. 2B), and screening results of turquoise color module genes under the filtering conditions of ‘geneSigFilter > 0.5 and moduleSigFilter > 0.75’.

### 3.3 Screening and validation of characteristic genes

The ‘veen’ package in R software was utilized to identify the overlapping genes between the selected PCA differential genes and the PCA candidate genes identified through WGCNA analysis (see Figure 3A). The shared genes were then filtered using the LASSO algorithm, resulting in the identification of three distinct genes (see Figure 3B),include Myristoylated alanine-rich C-kinase substrate like 1 (MARCKSL1), transmembrane and tetratricopeptide repeat containing 4(TMTC4), and Tubulin tyrosine ligase like 12 (TTLL12). Examination of the expression levels of these candidate genes in normal and tumor tissues revealed a significant increase in MARCKSL1, TMTC4, and TTLL12 in tumor patients, both in the training and test sets, with statistical significance (P<0.05) (see Figure 4A, B, C for training set data; D, E, F for test set data). Furthermore, ROC curve analysis of these characteristic genes in both sets demonstrated an AUC greater than 0.80 (see Figure 5A, B, C for training set data; D, E, F for test set data).

**Figure 3:**
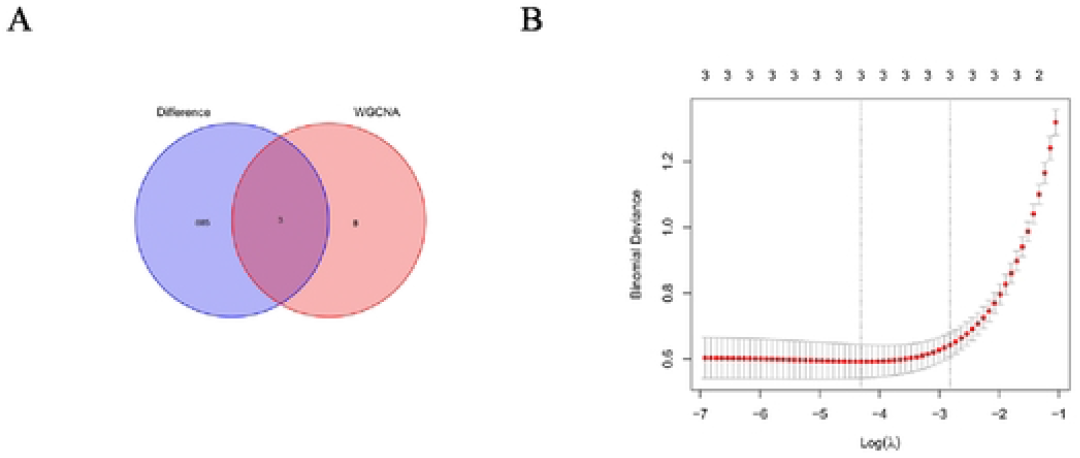
The veen diagram of the intersection of DEG and WNCGA (Figure 3A), LASSO analysis results (Figure 3B).

**Figure 4:**
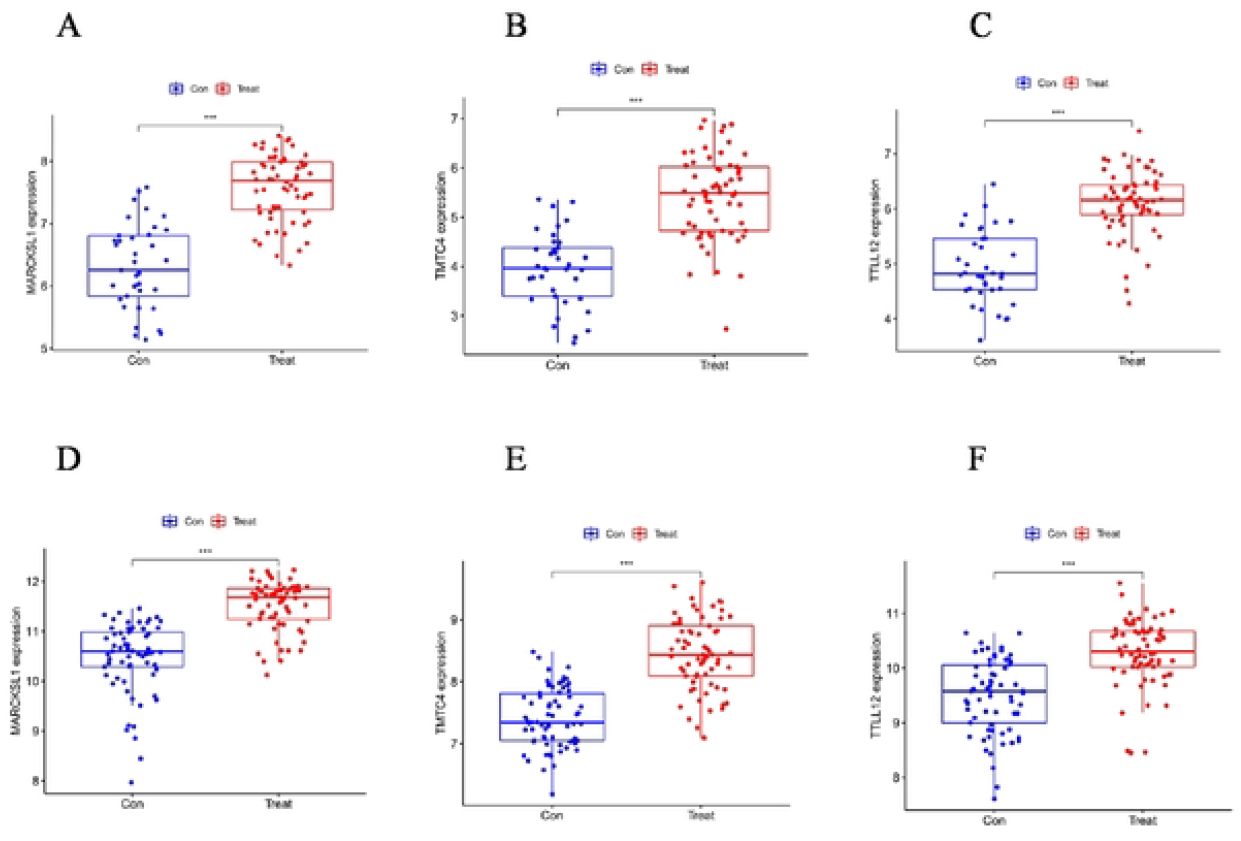
Differential expression analysis of candidate genes was conducted between normal individuals and PCA patients in both the training set (Fig.4A, B, C) and the test set (Fig.4D, E, F). In the comparison, normal individuals were considered as the control group while PCA patients were treated as the experimental group. Significance levels were denoted as follows: * for P < 0.05, ** for P < 0.01, and *** for P < 0.001.

**Figure 5:**
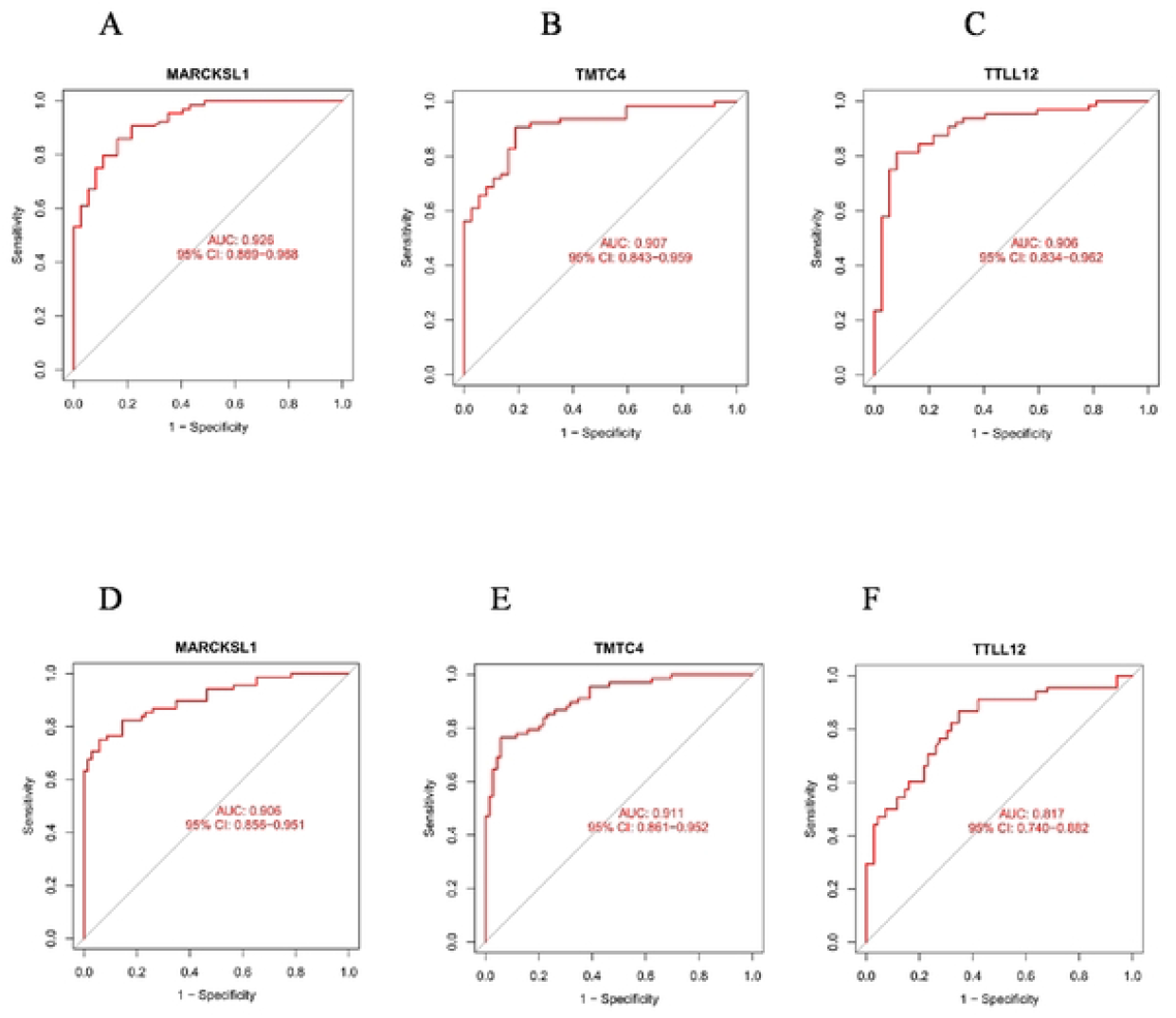
ROC analysis results of candidate genes in training set data samples (Figures 5A, B, C) and ROC analysis results of candidate genes in the test set data samples (Figures 5D, E, F).

### 3.4 Correlation analysis of key characteristic genes and immune cells

The immune correlation analysis was conducted on the characteristic genes with an AUC greater than 0.80 in both the training and validation sets, identifying Activated B cell, Eosinophil, Monocyte, Memory B cell, and Central memory CD4 T cell as infiltrated in tumor tissues. Conversely, CD56 dim natural killer cell, Immature dendritic cell, and Mast cell were found to be decreased in tumor tissues (Fig. 6A). MARCKSL1 showed a positive correlation with Plasmacytoid dendritic cell, Monocyte, and Central memory CD4 T cell (ρ < 0.05). TMTC4 exhibited a negative correlation with Natural killer T cell, Activated CD4 T cell, and Activated B cell, while showing a positive correlation with Central memory CD8 T cell and Central memory CD4 T cell (ρ < 0.05). TTLL12 displayed a negative correlation with Type 1 T helper cell, T follicular helper cell, and Activated CD4 T cell, but a positive correlation with Central memory CD4 T cell (ρ < 0.05) (Figure 6B). In conclusion, the immune cells infiltrated in tumor tissues that are significantly associated with the expression of key genes are Activated B cell, Monocyte, and Central memory CD4 T cell.

**Figure 6:**
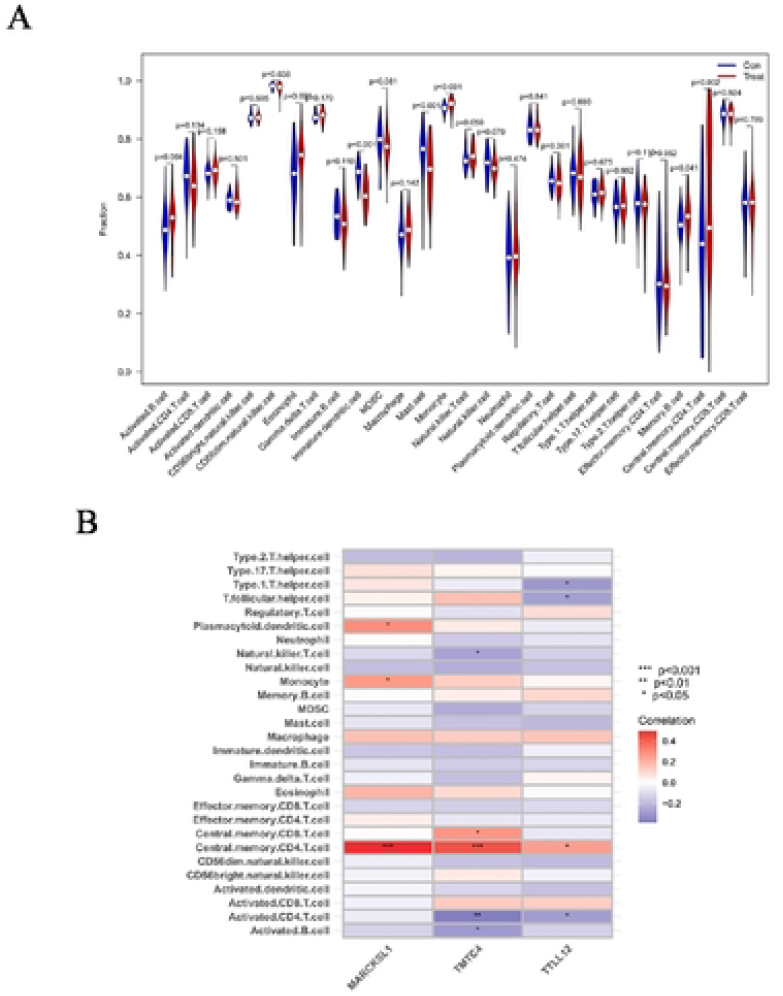
Comparison of immune cell infiltration in prostate tissue between individuals without prostate cancer (Con) and patients with prostate cancer (Treat) was conducted. A significance level of P < 0.05 was used to determine statistically significant differences (Figure 6A). The correlation between immune cells and the expression of key characteristic genes was analyzed, with red expression showing a positive correlation and dark expression showing a negative correlation. Significance levels were denoted as follows: • : P <0.05 ; ** : P <0.01 ; *** : P <0.001 (Figure 6B).

## 4. DISCUSSION

The occurrence and progression of PCA can be elusive, with early clinical symptoms often resembling those of benign prostatic hyperplasia. Currently, prostate specific antigen (PSA) is a common marker for early PCA diagnosis, but its reliability is affected by various factors like sexual activity and urinary tract infections. While PSA has strong specificity, it is known for its instability, high false positive rates, and limited utility in guiding treatment decisions^[6-7]^. In recent years, the relationship between genetic factors and diseases has garnered increasing attention among researchers. Genetic testing has gained popularity in clinical settings due to its stability, specificity, and effectiveness in diagnosis and treatment. Research indicates that PCA development is frequently associated with abnormal expression of oncogenes, yet further investigation in this area is warranted^[8-9]^. Therefore, identifying key characteristic genes involved in PCA development holds significant practical importance.

### 4.1 Key characteristic genes

In this study, the key characteristic genes of PCA were MARCKSL1, TMTC4 and TTLL12.

#### 4.1.1 MARCKSL1

MARCKSL1 belongs to the MARCKS family, also referred to as MARCKS-related protein (MRP), MARCKS-like protein (MLP), and brain protein F52^[10]^. MARCKSL1 is primarily localized on the cytoplasmic side of the membrane. Research indicates that MARCKSL1 undergoes phosphorylation by protein kinases, leading to its translocation from the cytoplasmic side of the membrane to the cytoplasm. It is selectively hydrolyzed by phospholipase C (PLC) upon interaction with phosphatidylinositol 4,5-bisphosphate (PIP2), resulting in the production of inositol triphosphate (IP3) and diacylglycerol (DAG). MARCKSL1 can also be phosphorylated by phosphoinositide 3-kinase (PI3K), leading to the generation of phosphatidylinositol 3,4,5-trisphosphate (PIP3). The phosphorylation status of MARCKSL1 influences cellular signaling pathways’ effectors, such as phospholipase D (PLD) and PI3K. The latter activates AKT signaling by phosphorylating PIP2 to PIP3, thereby regulating downstream signaling molecules^[11]^. This protein plays a role in regulating cell invasion and migration, vascular transport, various signal transduction processes, and other physiological activities^[12]^.

#### 4.1.2 TMTC4

The exact function of TMTC4, which targets cadherin 4, is not fully understood. It is primarily located in the endoplasmic reticulum (ER) and is involved in protein folding and calcium regulation^[13]^. Research has shown that due to the significant stress state of cancer cells, they rely on adaptive responses to survive, including the endoplasmic reticulum stress response (ER stress response) and unfolded protein response (UPR)^[14]^. Consequently, active cancer cell tissues often exhibit increased TMTC4 expression, which influences the metabolic processes of cancer cells through these mechanisms. The abnormal expression of TMTC4 has not been identified in other types of malignant tumors, suggesting that TMTC4 could potentially serve as a highly specific and sensitive diagnostic and therapeutic target for PCA, in line with findings by Rania M et al^[15]^.

#### 4.1.3 TTLL12

TTLL12 is involved in the post-translational modification of microtubules, which are essential fibrous structures in eukaryotic cells polymerized by tubulin. This modification, primarily carried out by tubulin tyrosine ligase (TTL)^[16]^, involves the attachment of nitrotyrosine to tubulin. The formation of nitrotyrosine tubulin (N-tubulin), specifically α-tubulin nitrotyrosination, has been linked to cancer cell death^[17]^. TTLL12 plays a role in reducing the formation of N-tubulin, potentially allowing cancer cells to evade this mechanism of cell death.

### 4.2 Analysis of immune function

Immunological analysis detected infiltration of Activated B cells, Monocytes, and Central memory CD4 T cells in tumor tissues, which exhibited a strong correlation with the expression of key characteristic genes.

#### 4.2.1 Activated B cell

B cells in the PCA tumor microenvironment have a multifaceted role, actively responding by presenting antigens and facilitating T cell activation during immune surveillance. Studies have shown clonal expansion in lymph nodes of PCA patients, with a rise in memory B cells and activated B cells, indicating tumor-specific T cell-dependent responses^[18]^. Moreover, the observed negative correlation between activated B cells and characteristic gene expression suggests that B cells may be suppressed in the PCA tumor microenvironment, leading to immune evasion by tumor cells.

#### 4.2.2 Monocytes

This study highlighted the infiltration of monocytes in PCA tumor tissues and their positive correlation with the expression of characteristic genes, suggesting a potential tumor-promoting effect. Monocytes have been found to secrete monocyte chemoattractant protein-1 (MCP-1), also known as chemokine ligand 2 (CCL2). The high-affinity receptor for human MCP family members, including MCP-1, CCL7, and CCL13, is chemokine receptor 2 (CCR2). Signaling through CCL2/CCR2 has been shown to promote tumor progression by enhancing cancer cell proliferation and survival, inducing migration and invasion, as well as stimulating inflammation and angiogenesis^[19]^. Moreover, CCL2/CCR2 can drive tumor epithelial mesenchymal transition (EMT) and the expression of MMP2 and MMP9 to facilitate cancer cell invasion, and attract tumor-associated macrophages (TAMs) to promote cancer cell dissemination. Additionally, CCL2 recruitment of TAMs and myeloid-derived suppressor cells (MDSCs) can trigger angiogenesis and suppress immune-mediated cancer cell attack, aligning with the findings of this study^[20]^.

#### 4.2.3 Central memory CD4 T cells

In T cell subsets, memory T cells play a crucial role in maintaining a lasting immune response and upholding the body’s anti-tumor immune status^[21]^. These memory T cells can be categorized into central memory T cells (Tcm) and effector memory T cells (Tem) based on their phenotype and function. Specifically, Tcm, characterized by high expression of CD45RO and CD62L, exhibit enhanced proliferation and anti-tumor capabilities compared to Tem^[22-23]^. Our study identified the infiltration of Tcm in the PCA tumor microenvironment, showing a positive correlation with the expression of characteristic genes. This suggests the presence of pre-existing immune memory in PCA patients, presenting as an immunosuppressive state. The potential mechanism and significance of elevated Tcm expression in the PCA tumor microenvironment include:➀The stimulation of memory T cells by tumor antigens leading to Tcm differentiation, immune memory formation, and increased inhibitory T cell expression, thereby suppressing the body’s anti-tumor immunity^[24]^**;➁**The migration of memory T cells to lymph nodes, where tumor invasion triggers continuous release of tumor antigens, promoting immune memory formation and Tcm differentiation, consequently reducing cytotoxic T lymphocytes (CTL) and diminishing their tumor cell killing efficacy^[25]^. These factors collectively contribute to immune evasion by PCA tumor cells, facilitating tumor progression.

## 4 CONCLUSIONS

The study utilized bioinformatics methods to identify key characteristic genes of PCA and examine their correlation with immune cells. This integration of bioinformatics and medicine using medical big data represents an interdisciplinary approach, offering a novel research methodology for scholars in various fields. However, the study is limited by the small sample size of PCA cases and the absence of experimental validation, resulting in a lack of in-depth understanding of the role of key characteristic genes in PCA development. To address this limitation, future research should focus on gathering larger sample data and conducting both basic and clinical studies to elucidate the mechanisms underlying the key genes in PCA progression.

## Declarations

### 1. Ethics Review

This research data was sourced from the GEO database and does not involve any animal or human experiments. As a result, it does not require review. Any issues that may arise will be the responsibility of the author.

### 2. Comments By The Unit

Agree to publish(Zhaotong Hospital of Traditional Chinese Medicine, Yunnan Province, China).

### 3. Availability of Data And Materials

The data involved in this study are all from the GEO database, and all data are open access.

### 5. Source of Funds

Project Fund : Yunnan University Joint Fund (No. : XYLH202352)

### 6. Author Contribution

Bo Shao : responsible for writing papers ; Pingping Sun: responsible for thesis revision and quality control ; Kaixiu Wu and Shui Wan: responsible for data analysis ; Yanggen Zuo, Li Xiao, Jinbo Pi, Zhengkai Fan, Zhongxiong Han are responsible for data collection.

### 7. Statement of thanks

Thank you to all the authors who have worked hard for this research institute. For your work, we are indispensable. We hope that our cooperation can continue forever and contribute to the development of medical undertakings.

